# A statistical framework to assess cross-frequency coupling while accounting for modeled confounding effects

**DOI:** 10.1101/519470

**Authors:** Jessica Nadalin, Louis-Emmanuel Martinet, Ethan Blackwood, Meng-Chen Lo, Alik S. Widge, Sydney S. Cash, Uri T. Eden, Mark A. Kramer

**Affiliations:** Department of Mathematics and Statistics, Boston University, Boston, Massachusetts 02215, USA; Department of Neurology, Massachusetts General Hospital, Boston, MA, 02114, USA; Department of Psychiatry, University of Minnesota, Minneapolis, Minnesota 55454, USA

## Abstract

Cross frequency coupling (CFC) is emerging as a fundamental feature of brain activity, correlated with brain function and dysfunction. Many different types of CFC have been identified through application of numerous data analysis methods, each developed to characterize a specific CFC type. Choosing an inappropriate method weakens statistical power and introduces opportunities for confounding effects. To address this, we propose a statistical modeling framework to estimate high frequency amplitude as a function of both the low frequency amplitude and low frequency phase; the result is a measure of phase-amplitude coupling that accounts for changes in the low frequency amplitude. We show in simulations that the proposed method successfully detects CFC between the low frequency phase or amplitude and the high frequency amplitude, and outperforms an existing method in biologically-motivated examples. Applying the method to *in vivo* data, we illustrate how CFC evolves during a seizure and is affected by electrical stimuli.

## Introduction

Brain rhythms - as recorded in the local field potential (LFP) or scalp electroencephalogram (EEG) - are believed to play a critical role in coordinating brain networks. By modulating neural excitability, these rhythmic fluctuations provide an effective means to control the timing of neuronal firing [23, 9]. Oscillatory rhythms have been categorized into different frequency bands (e.g., theta [4-10 Hz], gamma [30-80 Hz]) and associated with many functions: the theta band with memory, plasticity, and navigation [23]; the gamma band with local coupling and competition [39, 6]. In addition, gamma and high-gamma (80-200 Hz) activity have been identified as surrogate markers of neuronal firing [58, 50, 27, 56, 83, 61], observable in the EEG and LFP.

In general, lower frequency rhythms engage larger brain areas and modulate spatially localized fast activity [7, 14, 78, 44, 43]. For example, the phase of low frequency rhythms has been shown to modulate and coordinate neural spiking [77, 31, 26] via local circuit mechanisms that provide discrete windows of increased excitability. This interaction, in which fast activity is coupled to slower rhythms, is a common type of cross-frequency coupling (CFC). This particular type of CFC has been shown to carry behaviorally relevant information (e.g., related to position [32, 1], memory [65], decision making and coordination [20, 55, 87, 29]). More generally, CFC has been observed in many brain areas [7, 14, 19, 72, 24, 11], and linked to specific circuit and dynamical mechanisms [31]. The degree of CFC in those areas has been linked to working memory, neuronal computation, communication, learning and emotion [71, 33, 12, 21, 38, 46, 37, 36, 66], and clinical disorders [28, 86, 80, 4, 22], including epilepsy [82]. Although the cellular mechanisms giving rise to some neural rhythms are relatively well understood (e.g. gamma [85, 84, 48], and theta [71]), the neuronal substrate of CFC itself remains obscure.

Analysis of CFC focuses on relationships between the amplitude, phase, and frequency of two rhythms from different frequency bands. The notion of CFC, therefore, subsumes more specific types of coupling, including: phase-phase coupling (PPC), phase-amplitude coupling (PAC), and amplitude-amplitude coupling (AAC) [31]. PAC has been observed in rodent striatum and hippocampus [72] and human cortex [11], AAC has been observed between the alpha and gamma rhythms in dorsal and ventral cortices [57], and between theta and gamma rhythms during spatial navigation [64], and both PAC and AAC have been observed between alpha and gamma rhythms [53]. Many quantitative measures exist to characterize different types of CFC, including: mean vector length or modulation index [11, 70], phase-locking value [24, 42, 75], envelope-to-signal correlation [8], analysis of amplitude spectra [15], coherence between amplitude and signal [18], coherence between the time course of power and signal [53], and eigendecomposition of multichannel covariance matrices [16]. Overall, these different measures have been developed from different principles and made suitable for different purposes, as shown in comparative studies [70, 15, 54, 51].

Despite the richness of this methodological toolbox, it has limitations. For example, because each method focuses on one type of CFC, the choice of method restricts the type of CFC detectable in data. Applying a method to detect PAC in data with both PAC and AAC may: (i) falsely report no PAC in the data, or (ii) miss the presence of significant AAC in the same data. Changes in the low frequency power can also affect measures of PAC; increases in low frequency power can increase the signal to noise ratio of phase and amplitude variables, increasing the measure of PAC, even when the phase-amplitude coupling remains constant [3, 74, 33]. Furthermore, many experimental or clinical factors (e.g., stimulation parameters, age or sex of subject) can impact CFC in ways that are difficult to characterize with existing methods [17]. These observations suggest that an accurate measure of PAC would control for confounding variables, including the power of low frequency oscillations.

To that end, we propose here a generalized linear model (GLM) framework to assess CFC between the high-frequency amplitude and, simultaneously, the low frequency phase and amplitude. This formal statistical inference framework builds upon previous work [40, 54, 79, 74] to address the limitations of existing CFC measures. In what follows, we show that this framework successfully detects CFC in simulated signals. We compare this method to the modulation index, and show that in signals with CFC dependent on the low-frequency amplitude, the proposed method more accurately detects PAC than the modulation index. We apply this framework to *in vivo* recordings from human and rodent cortex and show examples of how accounting for AAC reveals changes in PAC over the course of seizure, and how to incorporate new covariates directly into the model framework.

## Methods

### Estimation of the phase and amplitude envelope

To study CFC we estimate three quantities: the phase of the low frequency signal, *ϕ*_low_; the amplitude envelope of the high frequency signal, *A*_high_; and the amplitude envelope of the low frequency signal, *A*_low_. To do so, we first bandpass filter the data into low frequency (4-7 Hz) and high frequency (100-140 Hz) signals, *V*_low_ and *V*_high_, respectively, using a least-squares linear-phase FIR filter of order 375 for the high frequency signal, and order 50 for the low frequency signal. Here we choose specific high and low frequency ranges of interest, motivated by previous *in vivo* observations [11, 72, 63]. However, we note that this method is flexible and not dependent on this choice, and that we select a wide high frequency band consistent with recommendations from the literature [3] and the mechanistic explanation that extracellular spikes produce this broadband high frequency activity [63]. We use the Hilbert transform to compute the analytic signals of *V*_low_ and *V*_high_, and from these compute the phase and amplitude of the low frequency signal (*A*_low_ and *ϕ*_low_) and the amplitude of the high frequency signal (*A*_high_).

### Modeling framework to assess CFC

Generalized linear models (GLMs) provide a principled framework to assess CFC [54, 40, 74]. Here, we present three models to analyze different types of CFC. The fundamental logic behind this approach is to model the distribution of *A*_high_ as a function of different predictors. In existing measures of PAC, the distribution of *A*_high_ versus *ϕ*_low_ is assessed using a variety of different metrics (e.g., [70]). Here, we estimate statistical models to fit *A*_high_ as a function of *ϕ*_low_, *A*_low_, and their combinations. If these models fit the data sufficiently well, then we estimate distances between the modeled surfaces to measure the impact of each predictor.

The *ϕ*_low_ model

The *ϕ*_*low*_ *model* relates *A*_high_, the response variable, to a linear combination of *ϕ*_low_, the predictor variable, expressed in a spline basis:

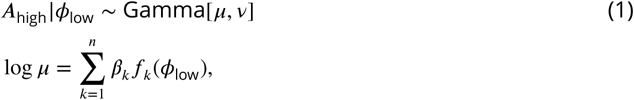

where the conditional distribution of *A*_high_ given *ϕ*_low_ is modeled as a Gamma random variable with mean parameter *µ* and shape parameter ν, and *β*_*k*_ are undetermined coefficients, which we refer to collectively as 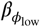 We choose this distribution as it guarantees real, positive amplitude values; we note that this distribution provides an acceptable fit to the example human data analyzed here (Figure 1). The functions {*f*_1_, …, *f*_*n*_} correspond to spline basis functions, with *n* control points equally spaced between 0 and 2*π*, used to approximate *ϕ*_low_. We use a tension parameter of 0.5, which controls the smoothness of the splines. We note that, because the link function of the conditional mean of the response variable (*A*_high_) varies linearly with the model coefficients *β*_*k*_ the model is a GLM. Here, we fix *n* = 10, which is a reasonable choice for smooth PAC with one or two broad peaks [40]. To support this choice, we apply an AIC-based selection procedure to 1000 simulated instances of signals of duration 20 s with phase-amplitude coupling and amplitude-amplitude coupling (see *Methods: Synthetic Time Series with PAC* and *Synthetic Time Series with AAC*, below, for simulation details). For each simulation, we fit Model 1 to these data for 27 different values of *n* from *n* = 4 to *n* = 30. For each simulated signal, we record the value of *n* such that we minimize the AIC, defined as

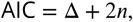

where Δ is the deviance from Model 1. The values of *n* that minimize the AIC tend to lie between *n* = 7 and *n* = 12 (Figure 2). These simulations support the choice of *n* = 10 as a sufficient number of splines.

**Figure 1.**
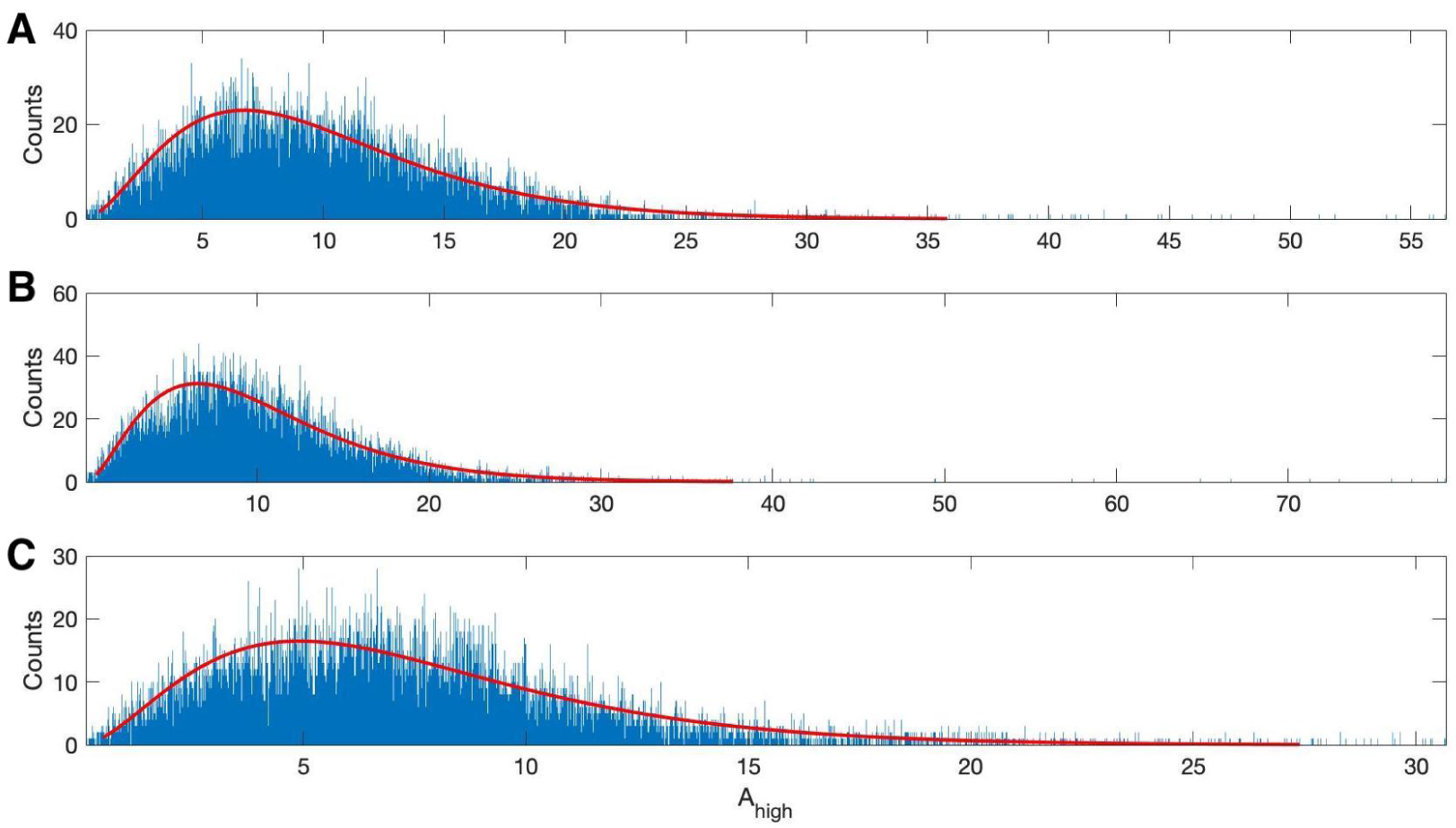
The gamma distribution provides a good fit to example human data. Three examples of 20s duration recorded from a single electrode during a human seizure. In each case, the gamma fit (red curve) provides an acceptable fit to the empirical distributions of the high frequency amplitude.

**Figure 2.**
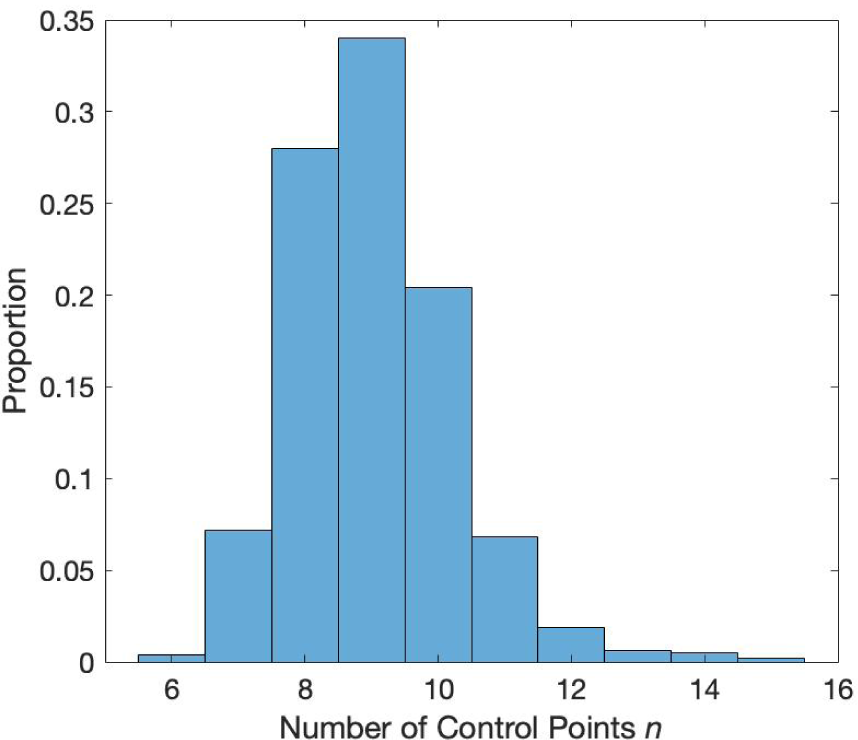
Distribution of the number of control points (*n*) that minimize the AIC. Values of *n* between 7 and 12 minimize the AIC in a simulation with phase-amplitude coupling and amplitude-amplitude coupling.

For a more detailed discussion and simulation examples of the PAC model, see [40]. We note that the choices of distribution and link function differ from those in [54, 74], where the normal distribution and identity link are used instead.

The *A*_low_ model

The *A*_*low*_ *model* relates the high frequency amplitude to the low frequency amplitude:

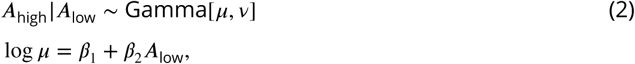

where the conditional distribution of *A*_high_ given *A*_low_ is modeled as a Gamma random variable with mean parameter *µ* and shape parameter ν. The predictor consists of a single variable and a constant, and the length of the coefficient vector 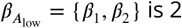 is 2.

The *A*_low_, *ϕ*_low_ model

The *A*_*low*_, *ϕ*_*low*_ *model* extends the *ϕ*_low_ model in Equation 1 by including three additional predictors in the GLM: *A*_low_, the low frequency amplitude; and interaction terms between the low frequency amplitude and the low frequency phase: *A*_low_ sin(*ϕ*_low_), and *A*_low_ cos(*ϕ*_low_). These new terms allow assessment of phase-amplitude coupling while accounting for linear amplitude-amplitude dependence and more complicated phase-dependent relationships on the low frequency amplitude without introducing many more parameters. Compared to the original *ϕ*_low_ model in Equation 1, including these new terms increases the number of variables to *n* + 3, and the length of the coefficient vector 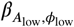 to *n* + 3. These changes result in the following model:

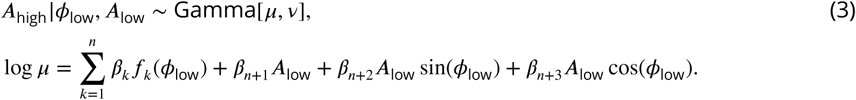

Here, the conditional distribution of *A*_high_ given *ϕ*_low_ and *A*_low_ is modeled as a Gamma random variable with mean parameter *µ* and shape parameter *ν*, and *β*_*k*_ are undetermined coefficients. We note that we only consider two interaction terms, rather than the spline basis function of phase, to reduce the number of parameters in the model.

### The statistics R_PAC_ and R_AAC_

We compute two measures of CFC, **R**_PAC_ and **R**_AAC_ which use the three models defined in the previous section. We evaluate each model in the three-dimensional space (*ϕ*_low_, *A*_low_, *A*_high_) and calculate the statistics **R**_PAC_ and **R**_AAC_. We use the MATLAB function *fitglm* to estimate the models; we note that this procedure estimates the dispersion directly for the gamma distribution. In what follows, we first discuss the three model surfaces estimated from the data, and then how we use these surfaces to compute the statistics **R**_PAC_ and **R**_AAC_.

To create the surface 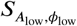, which fits the *A*_low_, *ϕ*_low_ model in the three-dimensional (*A*_low_, *ϕ*_low_, *A*_high_) space, we first compute estimates of the parameters 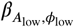 in Equation 3. We then estimate *A*_high_ by fixing *A*_low_ at one of 640 evenly spaced values between the 5th and 95th quantiles of *A*_low_ observed; we choose these quantiles to avoid extremely small or large values of *A*_low_. Finally, at the fixed *A*_low_, we compute the high frequency amplitude values from the *A*_low_, *ϕ*_low_ model over 100 evenly spaced values of *ϕ*_low_ between −*π* and *π*. This results in a two-dimensional curve 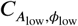 in the two-dimensional (*ϕ*_low_, *A*_high_) space with fixed *A*_low_. We repeat this procedure for all 640 values of *A*_low_ to create a surface 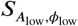 in the three-dimensional space (*A*_low_, *ϕ*_low_, *A*_high_) (Figure 3C).

**Figure 3.**
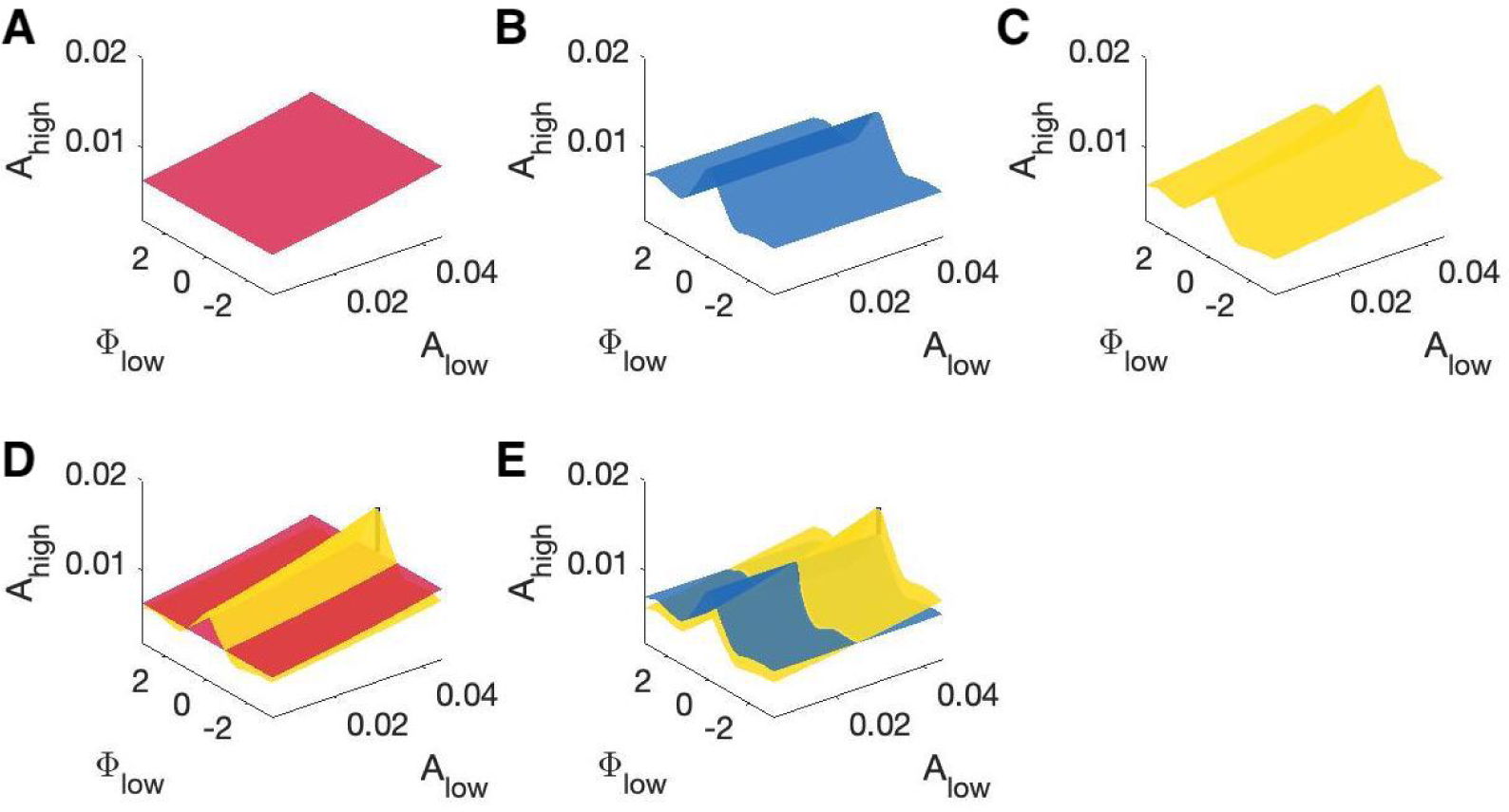
Example model surfaces used to determine R_PAC_ and R_AAC_. **(A,B,C)** Three example surfaces **(A)** 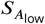, **(B)** 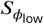, and **(C)** 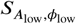 in the three-dimensional space (*A*_low_, *ϕ*_low_, *A*_high_). **(D)** The maximal distance between the surfaces 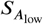 (red) and 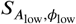 (yellow) is used to compute **R**_PAC_. **(E)** The maximal distance between the surfaces 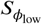 (blue) and 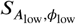 (yellow) is used to compute **R**_AAC_.

To create the surface 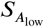, which fits the *A*_low_ model in the three-dimensional (*A*_low_, *ϕ*_low_, *A*_high_) space, we estimate the coefficient vector 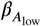 for the model in Equation 2. We then estimate the high frequency amplitude over 640 evenly spaced values between the 5th and 95th quantiles of *A*_low_ observed, again to avoid extremely small or large values of *A*_low_. This creates a mean response function which appears as a curve 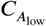 in the two-dimensional (*A*_low_, *A*_high_) space. We extend this two-dimensional curve to a three-dimensional surface 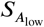, in the (*ϕ*_low_, *A*_high_) space, which extends 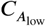 along the *ϕ*_low_ dimension (Figure 3A).

To create the surface 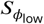, which fits the *ϕ*_low_ model in the three-dimensional (*A*_low_, *ϕ*_low_, *A*_high_) space, we first estimate the coefficients 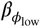 for the model in Equation 1. From this, we then compute estimates for the high frequency amplitude using the *ϕ*_low_ model with 100 evenly spaced values of *ϕ*_low_ between −*π* and *π*. Evaluating the *ϕ*_low_ model in this way results in a mean response function 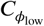 We extend this curve 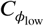 in the *A*_low_ dimension to create a surface 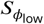 in the three-dimensional (*A*_low_, *ϕ*_low_, *A*_high_) space. The surface 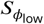 has the same structure as the curve 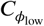 in the (*ϕ*_low_, *A*_high_) space, and remains constant along the dimension *A*_low_ (Figure 3B).

The statistic **R**_PAC_ measures the effect of low frequency phase on high frequency amplitude, while accounting for fluctuations in the low frequency amplitude. To compute this statistic, we note that the model in Equation 3 measures the combined effect of *A*_low_ and *ϕ*_low_ on *A*_high_, while the model in Equation 2 measures only the effect of *A*_low_ on *A*_high_. Hence, to isolate the effect of *ϕ*_low_ on *A*_high_, while accounting for *A*_low_, we compare the difference in fits between the models in Equations 2 and 3. We fit the mean response functions of the models in Equations 2 and 3, and calculate **R**_PAC_ as the maximum absolute fractional difference between the resulting surfaces 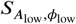 and 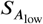 (Figure 3D):

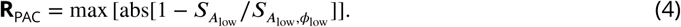

We expect fluctuations in 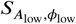 not present in 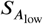 to be the result of *ϕ*_low_, i.e. PAC. In the absence of PAC, we expect the surfaces 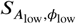 and 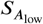 to be very close, resulting in a small value of **R**_PAC_. However, in the presence of PAC, we expect 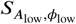 to deviate from 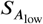, resulting in a large value of **R**_PAC_. We note that this measure, unlike R^2^ metrics for linear regression, is not meant to measure the goodness-of-fit of these models to the data, but rather the differences in fits between the two models. We also note that **R**_PAC_ is an unbounded measure, as it equals the maximum absolute fractional difference between distributions, which may exceed 1.

To compute the statistic **R**_AAC_, which measures the effect of low frequency amplitude on high frequency amplitude while accounting for fluctuations in the low frequency phase, we compare the difference in fits of the model in Equation 3 from the model in Equation 1. We note that the model in Equation 3 predicts *A*_high_ as a function of *A*_low_ and *ϕ*_low_, while the model in Equation 1 predicts *A*_high_ as a function of *ϕ*_low_ only. Therefore we expect a difference in fits between the models in Equations 1 and 3 results from the effects of *A*_low_ on *A*_high_. We fit the mean response functions of the models in Equations 1 and 3 in the three-dimensional (*ϕ*_low_, *A*_low_, *A*_high_) space, and calculate **R**_AAC_ as the maximum absolute fractional difference between the resulting surfaces 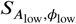 and 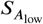 (Figure 3E):

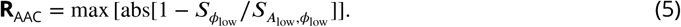

We expect fluctuations in 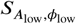 not present in 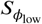 to be the result of *A*_low_, i.e. AAC. In the absence of AAC, we expect the surfaces 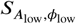 and 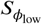 to be very close, resulting in a small value for **R**_AAC_. Alternatively, in the presence of AAC, we expect 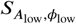 to deviate from 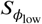 resulting in a large value of **R**_AAC_.

### Estimating 95% confidence intervals for R_PAC_ and R_AAC_

We compute 95% con1dence intervals for **R**_PAC_ and **R**_AAC_ via a parametric bootstrap method [40]. Given a vector of estimated coefficients *β*_x_ for *x* = {*A*_low_, *ϕ*_low_, or *A*_low_, *ϕ*_low_}, we use its estimated covariance and estimated mean to generate 10,000 normally distributed coefficient sample vectors 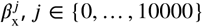. For each 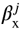, we then compute the high frequency amplitude values from the *A*_low_, *ϕ*_low_, or *A*_low_, *ϕ*_low_ model, 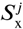. Finally, we compute the statistics 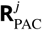 and 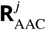 for each *j* as,

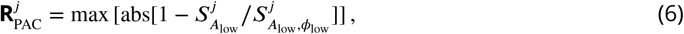

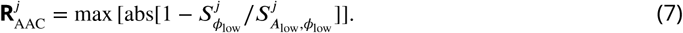

The 95% con1dence intervals for the statistics are the values of 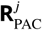 and 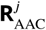 at the 0.025 and 0.975 quantiles [40].

### Assessing significance of AAC and PAC with bootstrap p-values

To assess whether evidence exists for significant PAC or AAC, we implement a bootstrap procedure to compute p-values as follows. Given two signals *V*_low_ and *V*_high_, and the resulting estimated statistics **R**_PAC_ and **R**_AAC_ we apply the Amplitude Adjusted Fourier Transform (AAFT) algorithm [69] on *V*_high_ to generate a surrogate signal 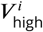. In the AAFT algorithm, we first reorder the values of *V*_high_ by creating a random Gaussian signal *W* and ordering the values of *V*_high_ to match *W*. For example, if the highest value of *W* occurs at index *j*, then the highest value of *V*_high_ will be reordered to occur at index *j*. Next, we apply the Fourier Transform (FT) to the reordered *V*_high_ and randomize the phase of the frequency domain signal. This signal is then inverse Fourier transformed and rescaled to have the same amplitude distribution as the original signal *V*_high_. In this way, the algorithm produces a permutation 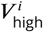 of *V*_high_ such that the power spectrum and amplitude distribution of the original signal are preserved.

We create 1000 such surrogate signals 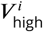, and calculate 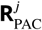 and 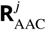 between *V*_low_ and each 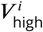. We define the p-values *p*_PAC_ and *p*_AAC_ as the proportion of values in 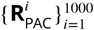 and 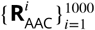 greater than the estimated statistics **R**_PAC_ and **R**_AAC_, respectively. If the proportion is zero, we set *p* = 0.0005.

We calculate p-values for the modulation index in the same way. The modulation index calculates the distribution of high frequency amplitudes versus low frequency phases and measures the distance from this distribution to a uniform distribution of amplitudes. Given the signals *V*_low_ and *V*_high_, and the resulting modulation index **MI** between them, we calculate the modulation index between *V*_low_ and 1000 surrogate permutations of *V*_high_ using the AAFT algorithm. We set *p*_MI_ to be the proportion of these resulting values greater than the **MI** value estimated from the original signals.

### Synthetic Time Series with PAC

We construct synthetic time series to examine the performance of the proposed method as follows. First, we simulate 20 s of pink noise data such that the power spectrum scales as 1/*f*. We then filter these data into low (4-7 Hz) and high (100-140 Hz) frequency bands, as described in *Methods: Estimation of the phase and amplitude envelope*, creating signals *V*_low_ and *V*_high_. Next, we couple the amplitude of the high frequency signal to the phase of the low frequency signal. To do so, we first locate the peaks of *V*_low_ and determine the times *t*_*k*_, *k* = {1, 2, 3, …, *K*}, of the *K* relative extrema. We note that these times correspond approximately to *ϕ*_low_ = 0. We then create a smooth modulation signal **M** which consists of a 42 ms Hanning window of height 1 + *I*_PAC_ centered at each *t*_*k*_, and a value of 1 at all other times (Figure 4A). The intensity parameter *I*_PAC_ in the modulation signal corresponds to the strength of PAC. *I*_PAC_ = 0.0 corresponds to no PAC, while *I*_PAC_ = 1.0 results in a 100% increase in the high frequency amplitude at each *t*_*k*_, creating strong PAC. We create a new signal 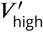 with the same phase as *V*_high_, but with amplitude dependent on the phase of *V*_low_ by setting,

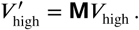

**Figure 4.**
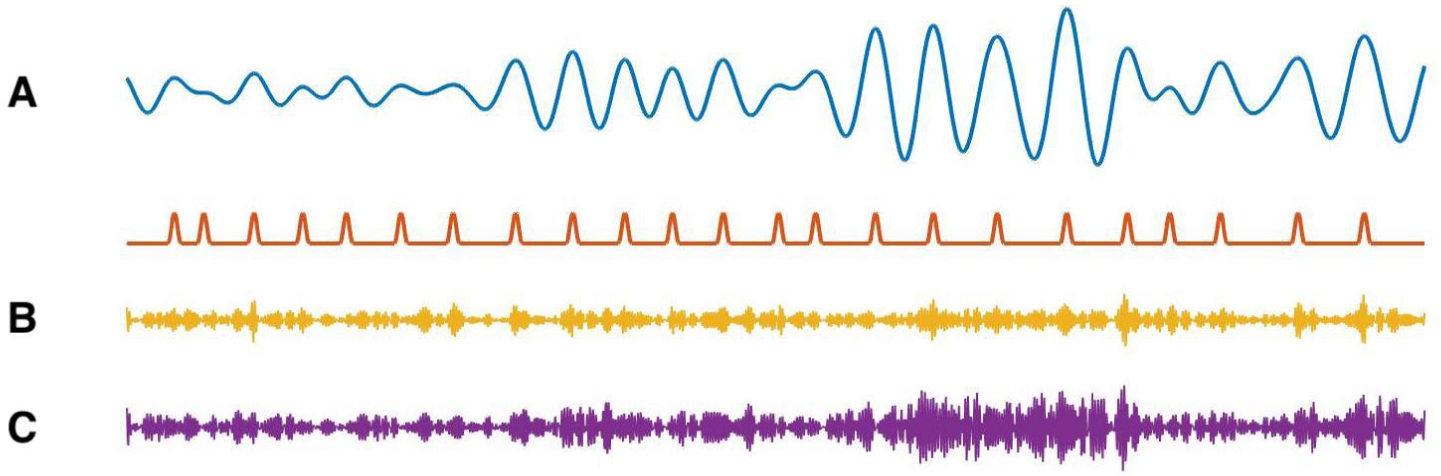
**(A)** Example simulation of *V*_low_ (blue) and modulation signal **M** (red). When the phase of *V*_low_ is 0 radians, **M** increases. **(B)** Example simulation of PAC. When the phase of *V*_low_ is approximately 0 radians, the high frequency amplitude (yellow) increases. **(C)** Example simulations of AAC. When the amplitude of *V*_low_ is large, so is the amplitude of the high frequency signal (purple).

We create the final voltage trace *V* as

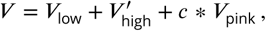

where *V*_pink_ is a new instance of pink noise multiplied by a small constant *c* = 0.01. In the signal *V*, brief increases of the high frequency activity occur at a specific phase (0 radians) of the low frequency signal (Figure 4B).

### Synthetic Time Series with AAC

To generate synthetic time series with dependence on the low frequency amplitude, we follow the procedure in the preceding section to generate *V*_low_, *V*_high_, and *A*_low_. We then induce amplitude-amplitude coupling between the low and high frequency components by creating a new signal 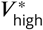 such that

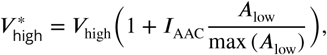

where *I*_AAC_ is the intensity parameter corresponding to the strength of amplitude-amplitude coupling. We define the final voltage trace *V* as

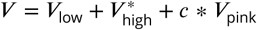

where *V*_pink_ is a new instance of pink noise multiplied by a small constant *c* = 0.01 (Figure 4C).

### Human Subject Data

A patient (male, age 32 years) with medically intractable focal epilepsy underwent clinically indicated intracranial cortical recordings for epilepsy monitoring. In addition to clinical electrode implantation, the patient was also implanted with a 10×10 (4 mm × 4 mm) NeuroPort microelectrode array (MEA; Blackrock Microsystems, Utah) in a neocortical area expected to be resected with high probability, in the temporal gyrus. The MEA consist of 96 platinum-tipped silicon probes, with a length of either 1-mm or 1.5-mm, corresponding to neocortical layer III as confirmed by histology after resection. Signals from the MEA were acquired continuously at 30 kHz per channel. Seizure onset times were determined by an experienced encephalographer (S.S.C.) through inspection of the macroelectrode recordings, referral to the clinical report, and clinical manifestations recorded on video. For a detailed clinical summary, see patient P2 of [81]. For these data, we analyze the 100-140 Hz frequency band to illustrate the proposed method; a more rigorous study of CFC in these data may require a more principled choice of high frequency band. This research was approved by local Institutional Review Boards at Massachusetts General Hospital and Brigham Women’s Hospitals (Partners Human Research Committee), and at Boston University according to National Institutes of Health guidelines.

### Code Availability

The code to perform this analysis is available for reuse and further development at https://github.com/Eden-Kramer-Lab/GLM-CFC.

## Results

We first examine the performance of the CFC measure through simulation examples. In doing so, we show that the statistics **R**_PAC_ and **R**_AAC_ accurately detect different types of cross-frequency coupling, increase with the intensity of coupling, and detect weak PAC coupled to the low frequency amplitude. We show that the proposed method is less sensitive to changes in low frequency power, and outperforms an existing PAC measure that lacks dependence on the low frequency amplitude. We conclude with example applications to human and rodent *in vivo* recordings, and propose that the new measure identifies cross-frequency coupling not detected in an existing PAC measure.

### The absence of CFC produces no significant detections of coupling

We first consider simulated signals without CFC. To create these signals, we follow the procedure in *Methods: Synthetic Time Series with PAC* with the modulation intensity set to zero (*I*_PAC_ = 0). In the resulting signals, *A*_high_ is approximately constant and does not depend on *ϕ*_low_ or *A*_low_ (Figure 5A). We estimate the *ϕ*_low_ model, the *A*_low_ model, and the *A*_low_, *ϕ*_low_ model from these data; we show example fits of the model surfaces in Figure 5B. We observe that the models exhibit small modulations in the estimated high frequency amplitude envelope as a function of the low frequency phase and amplitude.

**Figure 5.**
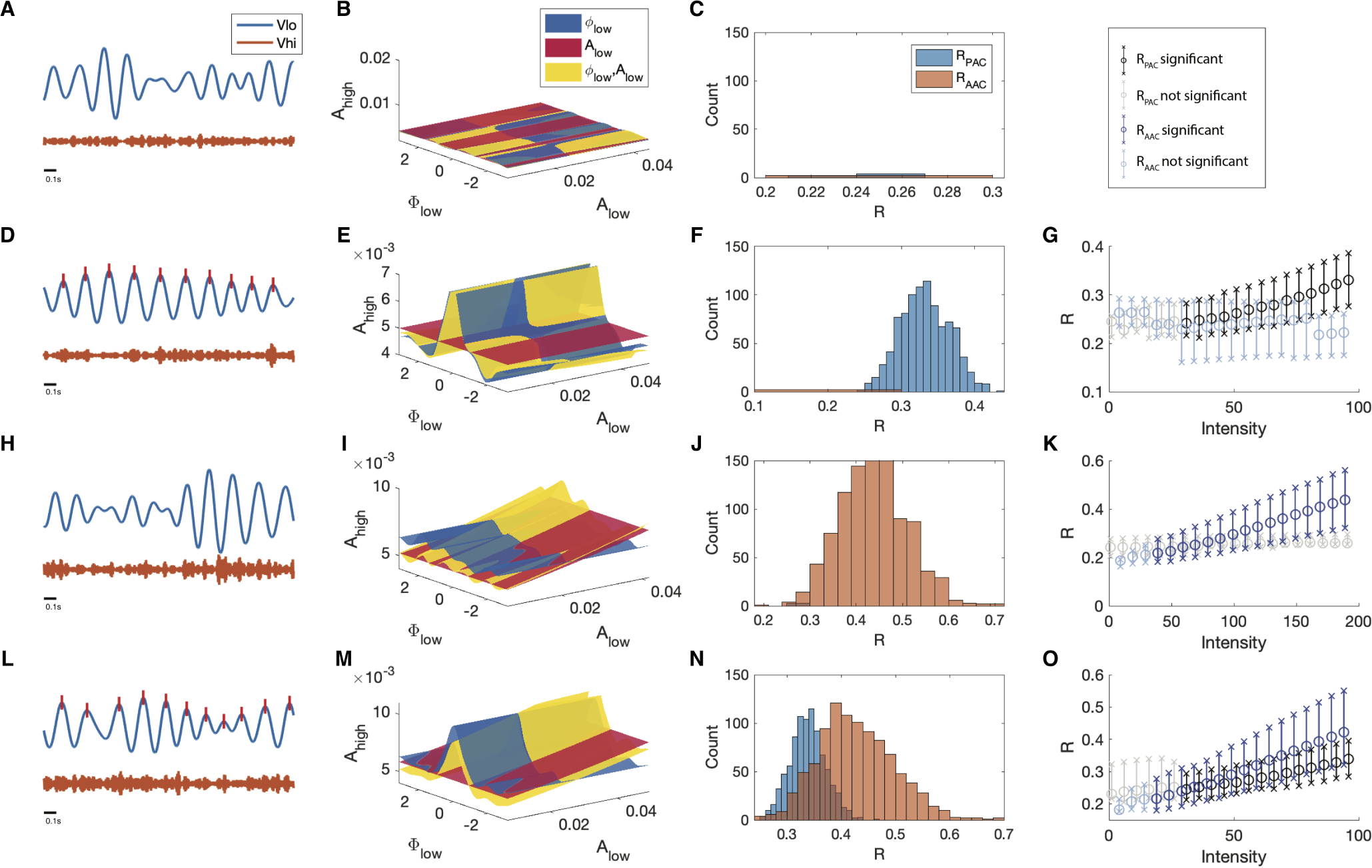
The statistical modeling framework successfully detects different types of cross-frequency coupling. **(A-C)** Simulations with no CFC. (A) When no CFC occurs, the low frequency signal (blue) and high frequency signal (orange) evolve independently. (B) The surfaces 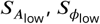, and 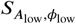 suggest no dependence of *A*_high_ on *ϕ*_low_ or *A*_low_. (C) Significant (*p*<0.05) values of **R**_PAC_ and **R**_AAC_ from 1000 simulations. Very few significant values for the statistics **R** are detected. **(D-G)** Simulations with PAC only. (D) When the phase of the low frequency signal is near 0 radians (red tick marks), the amplitude of the high frequency signal increases. (E) The surfaces 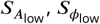 and 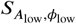 suggest dependence of *A*_high_ on *ϕ*_low_. (F) In 1000 simulations, significant values of **R**_PAC_ frequently appear, while significant values of **R**_AAC_ rarely appear. (G) As the intensity of PAC increases, so do the significant values of **R**_PAC_ (black), while any significant values of **R**_AAC_ remain small. **(H-K)** Simulations with AAC only. (H) The amplitudes of the high frequency signal and low frequency signal are positively correlated. (I) The surfaces 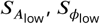, and 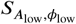 suggest dependence of *A*_high_ on *A*_low_. (J) In 1000 simulations, significant values of **R**_AAC_ frequently appear. (K) As the intensity of AAC increases, so do the significant values of **R**_AAC_ (blue), while any significant values of **R**_PAC_ remain small. **(L-O)** Simulations with PAC and AAC. (L) The amplitude of the high frequency signal increases when the phase of the low frequency signal is near 0 radians and the amplitude of the low frequency signal is large. (M) The surfaces 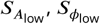, and 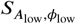 suggest dependence of *A*_high_ on *ϕ*_low_ and *A*_low_. (N) In 1000 simulations, significant values of **R**_PAC_ and **R**_AAC_ frequently appear. (O) As the intensity of PAC and AAC increase, so do the significant values of **R**_PAC_ and **R**_AAC_. In (G,K,O), circles indicate the median, and x’s the 5th and 95th quantiles.

To assess the distribution of significant **R** values in the case of no cross-frequency coupling, we simulate 1000 instances of the pink noise signals (each of 20 s) and apply the **R** measures to each instance, plotting significant **R** values in Figure 5C. We 1nd that for all 1000 instances, *p*_PAC_ and *p*_AAC_ are less than 0.05 in only 0.6% and 0.2% of the simulations, respectively, indicating no significant evidence of PAC or AAC, as expected.

### The proposed method accurately detects PAC

We next consider signals that possess phase-amplitude coupling, but lack amplitude-amplitude coupling. To do so, we simulate a 20 s signal with *A*_high_ modulated by *ϕ*_low_ (Figure 5D); more specifically, *A*_high_ increases when *ϕ*_low_ is near 0 radians (see *Methods, I*_PAC_ = 1). We then estimate the *ϕ*_low_ model, the *A*_low_ model, and the *A*_low_, *ϕ*_low_ model from these data; we show example fits in Figure 5E. We 1nd that the *ϕ*_low_ model is higher when *ϕ*_low_ is close to 0 radians, and the *A*_low_, *ϕ*_low_ model follows this trend. We note that, because the data do not depend on the low frequency amplitude (*A*_low_), the *ϕ*_low_ and *A*_low_, *ϕ*_low_ models have very similar shapes in the (*ϕ*_low_, *A*_low_, *A*_high_) space, and the *A*_low_ model is nearly flat.

Simulating 1000 instances of these 20 s signals with induced phase-amplitude coupling, we 1nd *p*_AAC_ *<* 0.05 for only 0.6% of the simulations, while *p*_PAC_ *<* 0.05 for 96.5% of the simulations. We 1nd that the significant values of **R**_PAC_ lie well above 0 (Figure 5F), and that as the intensity of the simulated phase-amplitude coupling increases, so does the statistic **R**_PAC_ (Figure 5G). We conclude that the proposed method accurately detects the presence of phase-amplitude coupling in these simulated data.

### The proposed method accurately detects AAC

We next consider signals with amplitude-amplitude coupling, but without phase-amplitude coupling. We simulate a 20 s signal such that *A*_high_ is modulated by *A*_low_ (see *Methods, I*_AAC_ = 1); when *A*_low_ is large, so is *A*_high_ (Figure 5H). We then estimate the *ϕ*_low_ model, the *A*_low_ model, and the *A*_low_, *ϕ*_low_ model (example fits in Figure 5I). We 1nd that the *A*_low_ model increases along the *A*_low_ axis, and that the *A*_low_, *ϕ*_low_ model closely follows this trend, while the *ϕ*_low_ model remains mostly flat, as expected.

Simulating 1000 instances of these signals we 1nd that *p*_AAC_ *<* 0.05 for 97.9% of simulations, while *p*_PAC_ *<* 0.05 for 0.3% of simulations. The significant values of **R**_AAC_ lie above 0 (Figure 5J), and increases in the intensity of AAC produce increases in **R**_AAC_ (Figure 5K). We conclude that the proposed method accurately detects the presence of amplitude-amplitude coupling.

### The proposed method accurately detects the simultaneous occurrence of PAC and AAC

We now consider signals that possess both phase-amplitude coupling and amplitude-amplitude coupling. To do so, we simulate time series data with both AAC and PAC (Figure 5L). In this case, *A*_high_ increases when *ϕ*_low_ is near 0 radians and when *A*_low_ is large (see *Methods, I*_PAC_ = 1 and *I*_AAC_ = 1). We then estimate the *ϕ*_low_ model, the *A*_low_ model, and the *A*_low_, *ϕ*_low_ model from the data and visualize the results (Figure 5M). We find that the *ϕ*_low_ model increases near *ϕ*_low_ = 0, and that the *A*_low_ model increases linearly with *A*_low_. The *A*_low_, *ϕ*_low_ model exhibits both of these behaviors, increasing at *ϕ*_low_ = 0 and as *A*_low_ increases.

Simulating 1000 instances of signals with both AAC and PAC present, we find that *p*_AAC_ < 0.05 in 96.7% of simulations and *p*_PAC_ < 0.05 in 98.1% of simulations. The distributions of significant **R**_PAC_ and **R**_AAC_ values lie above 0, consistent with the presence of both PAC and AAC (Figure 5N), and as the intensity of PAC and AAC increases, so do the values of **R**_PAC_ and **R**_AAC_ (Figure 5O). We conclude that the model successfully detects the concurrent presence of PAC and AAC.

### R_PAC_ and modulation index are both sensitive to weak modulations

To investigate the ability of the proposed method and the modulation index to detect weak coupling between the low frequency phase and high frequency amplitude, we perform the following simulations. For each intensity value *I*_PAC_ between 0 and 0.5 (in steps of 0.025), we simulate 1000 signals (see *Methods*) and compute **R**_PAC_ and a measure of PAC in common use: the modulation index **MI** [70] (Figure 6). We find that both **MI** and **R**_PAC_, while small, increase with *I*_PAC_; in this way, both measures are sensitive to small values of *I*_PAC_.

**Figure 6.**
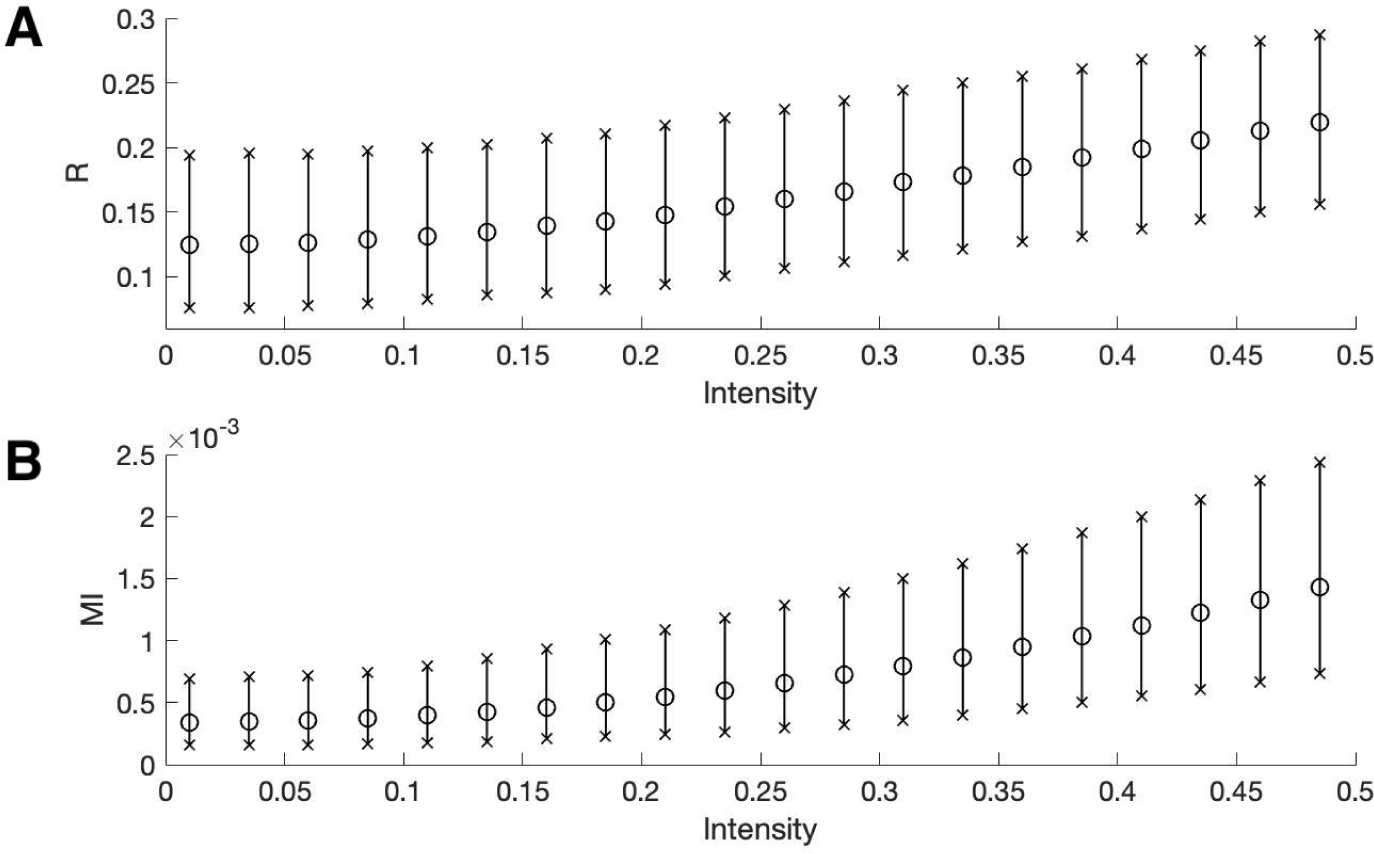
The two measures of PAC increase with intensities near zero. The mean (circles) and 5^th^ to 95^th^ quantiles (x’s) of **(A) R**_PAC_ and **(B) MI** for intensity values between 0 and 0.5. Both measures increase with the intensity.

### The proposed method is less affected by fluctuations in low-frequency amplitude and AAC

Increases in low frequency power can increase measures of phase-amplitude coupling, although the underlying PAC remains unchanged [3, 17]. Characterizing the impact of this confounding effect is important both to understand measure performance and to produce accurate interpretations of analyzed data. To examine this phenomenon, we perform the following simulation. First, we simulate a signal *V* with fixed PAC (intensity *I*_PAC_ = 1, see *Methods*). Second, we filter *V* into its low and high frequency components *V*_low_ and *V*_high_, respectively. Then, we create a new signal *V* * as follows:

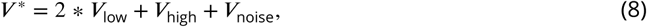

where *V*_noise_ is a pink noise term (see *Methods*). We note that we only alter the low frequency component of *V* and do not alter the PAC. To analyze the PAC in this new signal we compute **R**_PAC_ and **MI**.

We show in Figure 7 population results (1000 realizations each of the simulated signals *V* and *V* *) for the **R** and **MI** values. We observe that increases in the amplitude of *V*_low_ produce increases in **MI** and **R**_PAC_. However, this increase is more dramatic for **MI** than for **R**_PAC_; we note that the distributions of **R**_PAC_ almost completely overlap (Figure 7A), while the distribution of **MI** shifts to larger values when the amplitude of *V*_low_ increases (Figure 7B). We conclude that the statistic **R**_PAC_ — which includes the low frequency amplitude as a predictor in the GLM — is more robust to increases in low frequency power than a method that only includes the low frequency phase.

**Figure 7.**
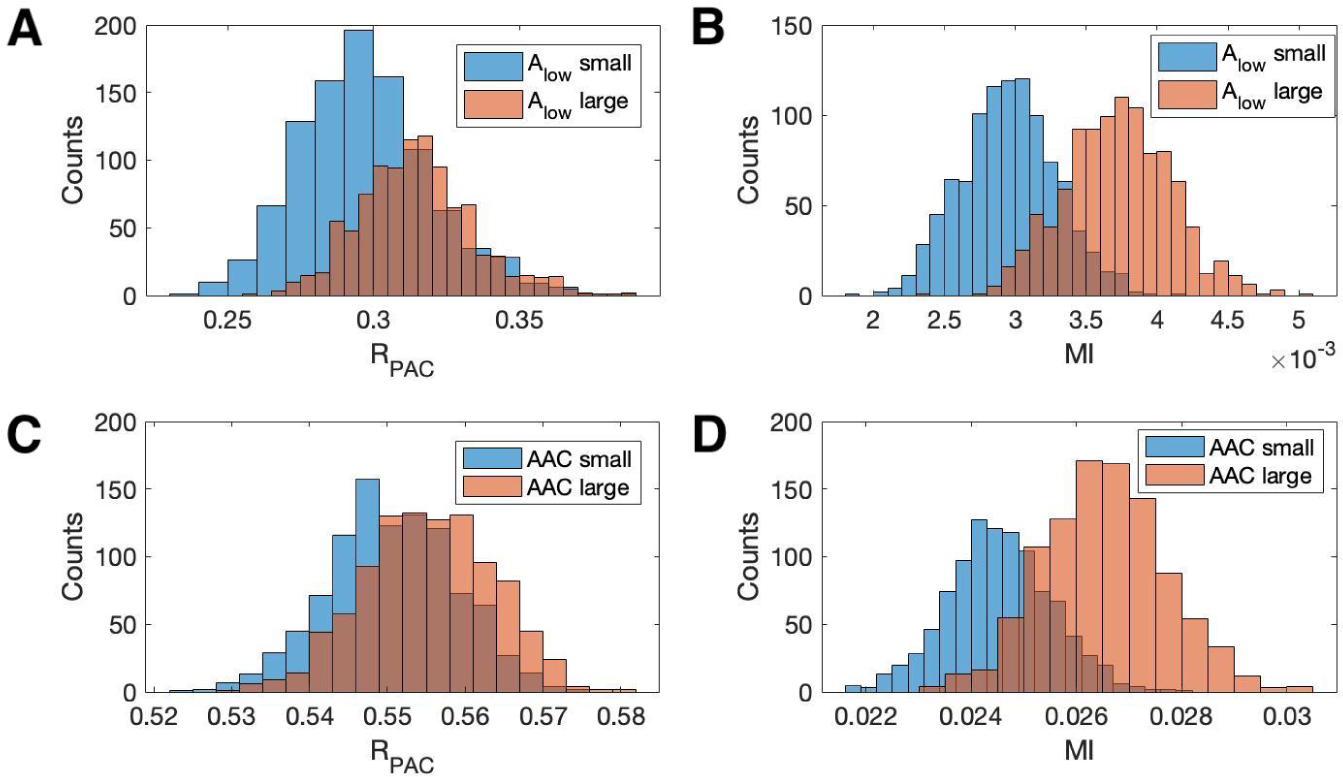
Increases in the amplitude of the low frequency signal, and the amplitude-amplitude coupling (AAC), increase the modulation index more than R_PAC_. **(A,B)** Distributions of (A) **R**_PAC_ and (B) **MI** when *A*_low_ is small (blue) and when *A*_low_ is large (red). **(C,D)** Distributions of (C) **R**_PAC_ and (D) **MI** when AAC is small (blue) and when AAC is large (red).

We also investigate the effect of increases in amplitude-amplitude coupling (AAC) on the two measures of PAC. As before, we simulate a signal *V* with fixed PAC (intensity *I*_PAC_ = 1) and no AAC (intensity *I*_AAC_ = 0). We then simulate a second signal *V*\* with the same fixed PAC as *V*, and with additional AAC (intensity *I*_AAC_ = 10). We simulate 1000 realizations of *V* and *V*\* and compute the corresponding **R**_PAC_ and **MI** values. We observe that the increase in AAC produces a small increase in the distribution of **R**_PAC_ values (Figure 7C), but a large increase in the distribution of **MI** values (Figure 7D). We conclude that the statistic **R**_PAC_ is more robust to increases in AAC than **MI**.

These simulations show that at a fixed, non-zero PAC, the modulation index increases with increased *A*_low_ and AAC. We now consider the scenario of increased *A*_low_ and AAC in the absence of PAC. To do so, we simulate 1000 signals of 200 s duration, with no PAC (intensity *I*_PAC_ = 0). For each signal, at time 100 s (i.e., the midpoint of the simulation) we increase the low frequency amplitude by a factor of 10 (consistent with observations from an experiment in rodent cortex, as described below), and include AAC between the low and high frequency signals (intensity *I*_AAC_ = 0 for *t* < 100 s and intensity *I*_AAC_ = 2 for *t* ≥ 100 s). We find that, in the absence of PAC, **R**_PAC_ detects significant PAC (p<0.05) in 0.4% of the simulated signals, while **MI** detects significant PAC in 34.3% of simulated signals. We conclude that in the presence of increased low frequency amplitude and amplitude-amplitude coupling, **MI** may detect PAC where none exists, while **R**_PAC_, which accounts for fluctuations in low frequency amplitude, does not.

### Sparse PAC is detected when coupled to the low frequency amplitude

While the modulation index has been successfully applied in many contexts [12, 31], instances may exist where this measure is not optimal. For example, because the modulation index was not designed to account for the low frequency amplitude, it may fail to detect PAC when *A*_high_ depends not only on *ϕ*_low_, but also on *A*_low_. For example, since the modulation index considers the distribution of *A*_high_ at all observed values of *ϕ*_low_, it may fail to detect coupling events that occur sparsely at only a subset of appropriate *ϕ*_low_ occurrences. **R**_PAC_, on the other hand, may detect these sparse events if these events are coupled to *A*_low_, as **R**_PAC_ accounts for fluctuations in low frequency amplitude. To illustrate this, we consider a simulation scenario in which PAC occurs sparsely in time.

We create a signal *V* with PAC, and corresponding modulation signal **M** with intensity value *I*_PAC_ = 1.0 (see *Methods*, Figure 8A-B). We then modify this signal to reduce the number of PAC events in a way that depends on *A*_low_. To do so, we preserve PAC at the peaks of *V*_low_ (i.e., when *ϕ*_low_ = 0), but now only when these peaks are large, more specifically in the top 5% of peak values.

**Figure 8.**
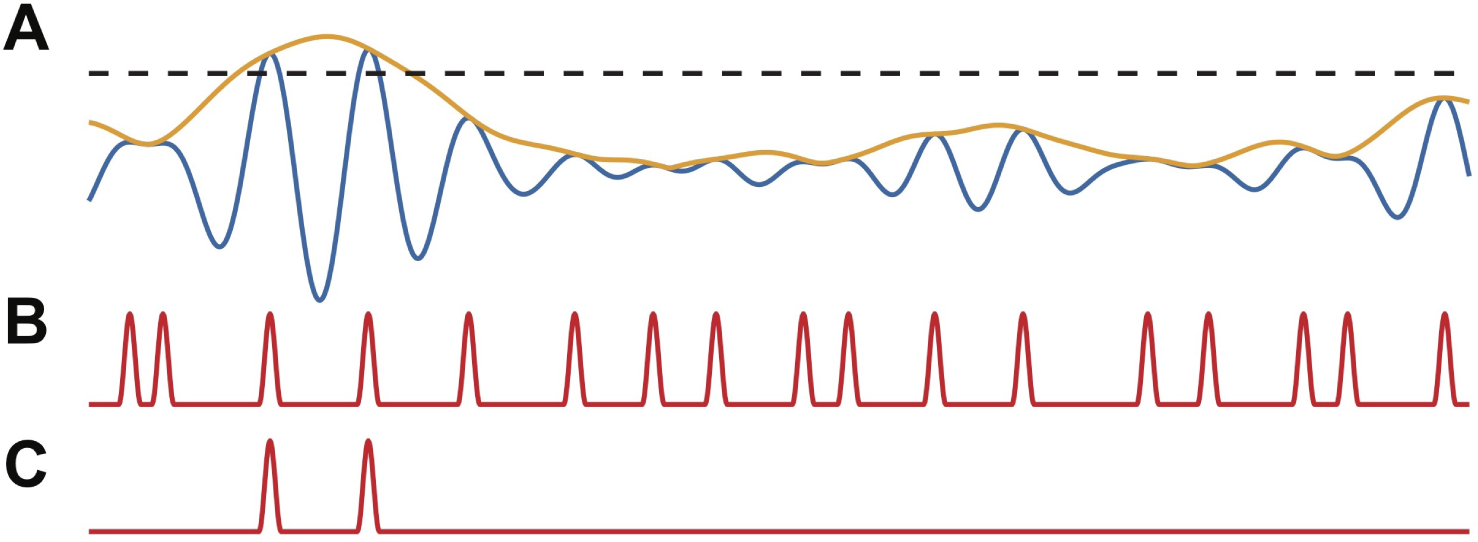
PAC events restricted to a subset of occurrences are still detectable. **(A)** The low frequency signal (blue), amplitude envelope (yellow), and threshold (black dashed). **(B-C)** The modulation signal increases (B) at every occurrence of *ϕ*_low_ = 0, or (C) only when *A*_low_ exceeds the threshold and *ϕ*_low_ = 0.

We define a threshold value *T* to be the 95^th^ quantile of the peak *V*_low_ values, and modify the modulation signal **M** as follows. When **M** exceeds 1 (i.e., when *ϕ*_low_ = 0) and the low frequency amplitude exceeds *T* (i.e., *A*_low_ ≥ *T*), we make no change to **M**. Alternatively, when **M** exceeds 1 and the low frequency amplitude lies below *T* (i.e., *A*_low_ < *T*), we decrease **M** to 1 (Figure 8C). In this way, we create a modified modulation signal **M**_1_ such that in the resulting signal *V*_1_, when *ϕ*_low_ = 0 and *A*_low_ is large enough, *A*_high_ is increased; and when *ϕ*_low_ = 0 and *A*_low_ is not large enough, there is no change to *A*_high_. This signal *V*_1_ hence has fewer phase-amplitude coupling events than the number of times *ϕ*_low_ = 0.

We generate 1000 realizations of the simulated signals *V*_1_, and compute **R**_PAC_ and **MI**. We find that while **MI** detects significant PAC in only 37% of simulations, **R**_PAC_ detects significant PAC in 72% of simulations. In this case, although the PAC occurs infrequently, these occurrences are coupled to *A*_low_, and **R**_PAC_, which accounts for changes in *A*_low_, successfully detects these events much more frequently. We conclude that when the PAC is dependent on *A*_low_, **R**_PAC_ more accurately detects these sparse coupling events.

### The CFC model detects simultaneous PAC and AAC missed in an existing method

To further illustrate the utility of the proposed method, we consider another scenario in which *A*_low_ impacts the occurrence of PAC. More specifically, we consider a case in which *A*_high_ increases at a fixed low frequency phase for high values of *A*_low_, and *A*_high_ decreases at the same phase for small values of *A*_low_. In this case, we expect that the modulation index may fail to detect the coupling because the distribution of *A*_high_ over *ϕ*_low_ would appear uniform when averaged over all values of *A*_low_; the dependence of *A*_high_ on *ϕ*_low_ would only become apparent after accounting for *A*_low_.

To implement this scenario, we consider the modulation signal **M** (see *Methods*) with an intensity value *I*_PAC_ = 1. We consider all peaks of *A*_low_ and set the threshold *T* to be the 50^th^ quantile (Figure 9A). We then modify the modulation signal **M** as follows. When **M** exceeds 1 (i.e., when *ϕ*_low_ = 0) and the low frequency amplitude exceeds *T* (i.e., *A*_low_ ≥ *T)*, we make no change to **M**. Alternatively, when **M** exceeds 1 and the low frequency amplitude lies below *T* (i.e. *A*_low_ < *T)*, we decrease **M** to 0 (Figure 9B). In this way, we create a modified modulation signal **M** such that when *ϕ*_low_ = 0 and *A*_low_ is large enough, *A*_high_ is increased; and when *ϕ*_low_ = 0 and *A*_low_ is small enough, *A*_high_ is decreased (Figure 9C).

**Figure 9.**
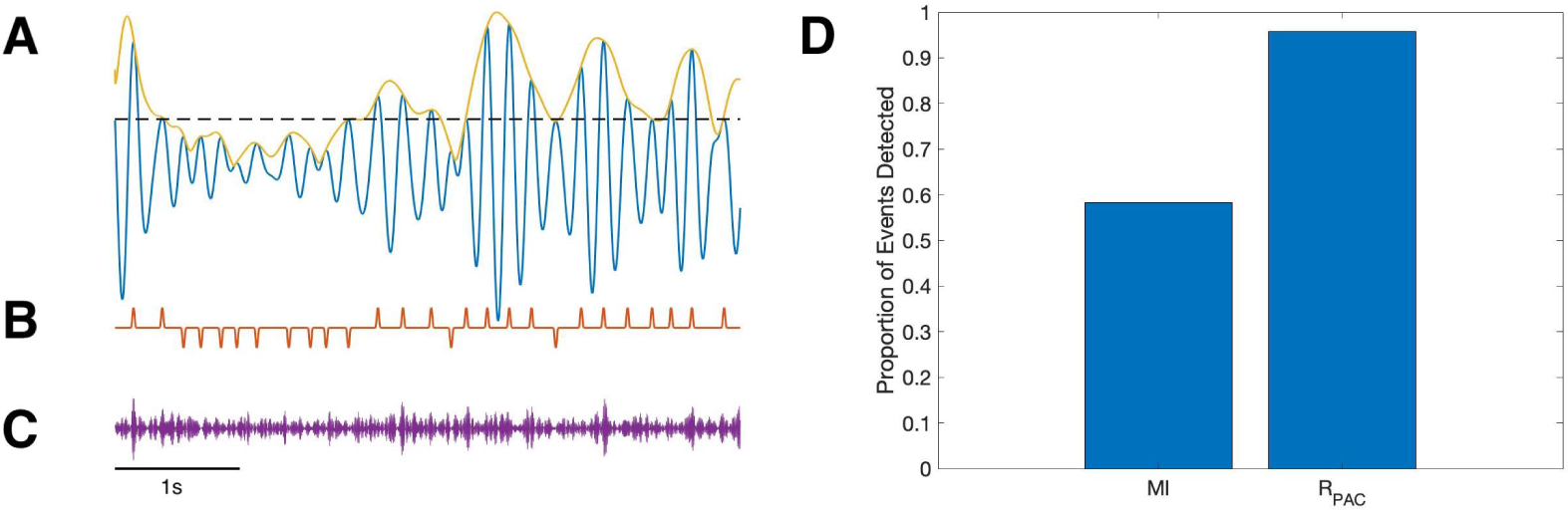
PAC with AAC is accurately detected with the proposed method, but not with the modulation index. **(A)** The low frequency signal (blue), amplitude envelope (yellow), and threshold (black dashed). **(B)** The modulation signal (red) increases when *ϕ*_low_ = 0 and *A*_low_ > *T*, and deceases when *ϕ*_low_ = 0 and *A*_low_ < *T*. **(C)** The modulated *A*_high_ signal (purple) increases and decreases with the modulation signal. **(D)** The proportion of significant detections (out of 1000) for **MI** and **R**_PAC_.

Using this method, we simulate 1000 realizations of this signal, and calculate **MI** and **R**_PAC_ for each signal (Figure 9D). We find that **R**_PAC_ detects significant PAC in nearly all (96%) of the simulations, while **MI** detects significant PAC in only 58% of the simulations. We conclude that, in this simulation, **R**_PAC_ more accurately detects PAC coupled to low frequency amplitude than the modulation index.

### A simple stochastic spiking neural model illustrates the utility of the proposed method

In the previous simulations, we created synthetic data without a biophysically principled generative model. Here we consider an alternative simulation strategy with a more direct connection to neural dynamics. While many biophysically motivated models of cross-frequency coupling exist [62, 13, 67, 30, 45, 52, 25, 47, 35, 68, 88, 73], we consider here a relatively simple stochastic spiking neuron model [2]. In this stochastic model, we generate a spike train (*V*_high_) in which *V*_low_ modulates the probability of spiking as a function of *A*_low_ and *ϕ*_low_. We note that high frequency activity is thought to represent the aggregate spiking activity of local neural populations [61, 10, 59, 34]; while here we simulate the activity of a single neuron, the spike train still produces temporally focal events of high frequency activity. In this framework, we allow the target phase 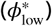 modulating *A*_high_ to change as a function of *A*_low_: when *A*_low_ is large, the probability of spiking is highest near *ϕ*_low_ = ±*π*, and when *A*_low_ is small, the probability of spiking is highest near *ϕ*_low_ = 0. More precisely, we define 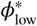 as

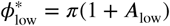

where *A*_low_ is a sinusoid oscillating between 1 and 2 with period 0.1 Hz. We define the spiking probability, *λ*, as

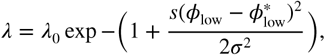

where (*σ* = 0.01, *s*(*ϕ*) is a triangle wave, and we choose *λ*_0_ so that the maximum value of *λ* is 2. We note that the spiking probability *λ* is zero except near times when the phase of the low frequency signal (*ϕ*_low_) is near 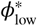. We then define *A*_high_ as:

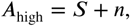

where *S* is the binary sequence generated by the stochastic spiking neuron model, and *n* is Gaussian noise with mean zero and standard deviation 0.1. In this scenario, the distribution of *A*_high_ over *ϕ*_low_ appears uniform when averaged over all values of *A*_low_. We therefore expect the modulation index to remain small, despite the presence of PAC with maximal phase dependent on *A*_low_. However, we expect that **R**_PAC_, which accounts for fluctuations in low frequency amplitude, will detect this PAC. We show an example signal from this simulation in Figure 10A. As expected, we find that **R**_PAC_ detects PAC (**R**_PAC_ = 0.172, *p* = 0.02); we note that the (*A*_low_, *ϕ*_low_) surface exhibits a single peak near *ϕ*_low_ = 0 at small values of *A*_low_, and at *ϕ*_low_ = ±*π* at large value of *A*_low_ (Figure 10B). The (*A*_low_, *ϕ*_low_) surface deviates significantly from the *A*_low_ surface, resulting in a large **R**_PAC_ value. However, the non-uniform shape of the (*A*_low_, *ϕ*_low_) surface is lost when we fail to account for *A*_low_. In this scenario, the distribution of *A*_high_ over *ϕ*_low_ appears uniform, resulting in a low **MI** value (Figure 10C).

**Figure 10.**
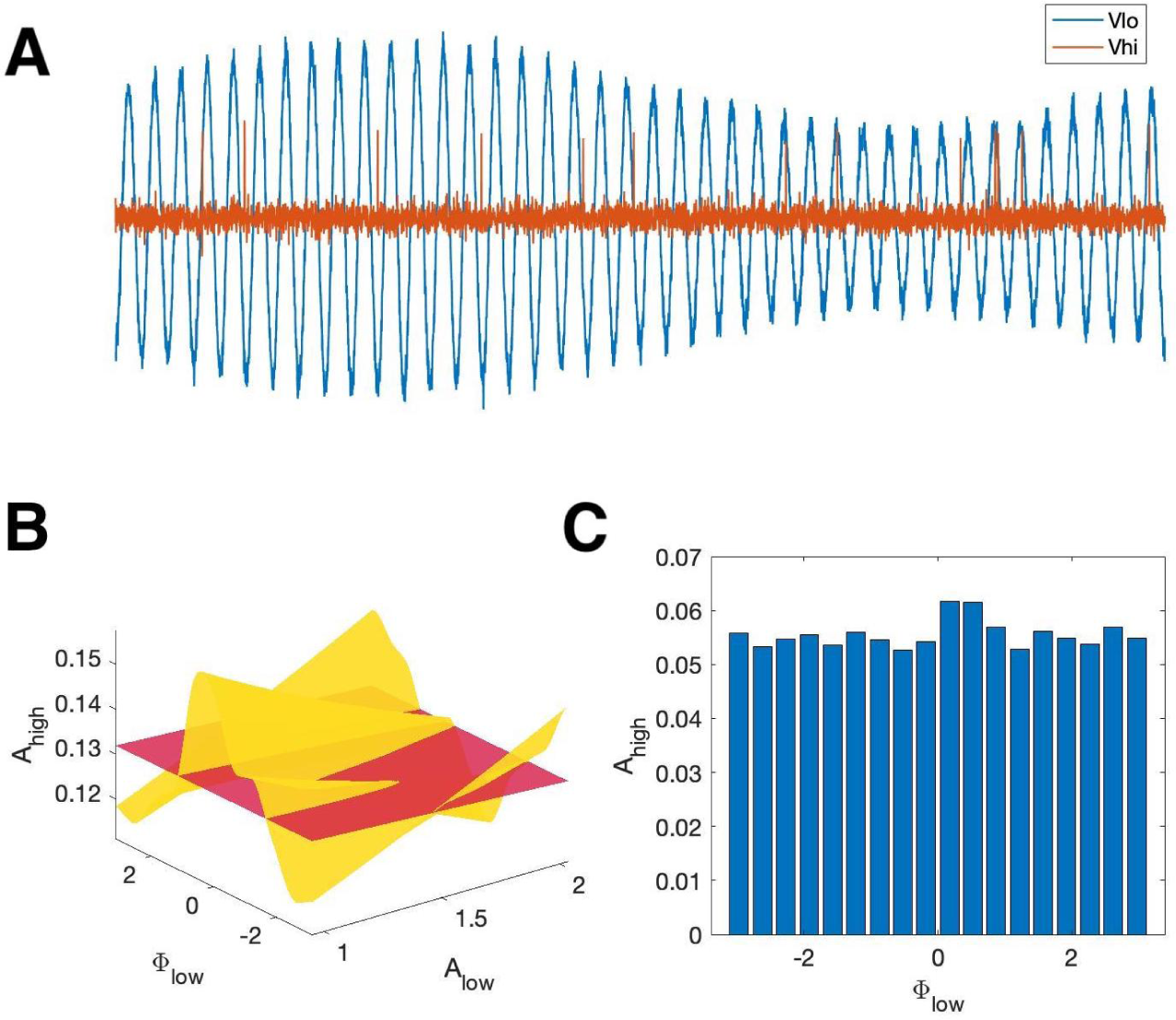
R_PAC_, but not MI, detects phase amplitude coupling in a simple stochastic spiking neuron model. **(A)** The phase and amplitude of the low frequency signal (blue) modulate the probability of a high frequency spike (orange). **(B)** The surfaces 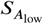 (red) and 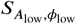 (yellow). The phase of maximal *A*_high_ modulation depends on *A*_low_. **(C)** The modulation index fails to detect this type of PAC.

### Application to *in vivo* human seizure data

To evaluate the performance of the proposed method on *in vivo* data, we first consider an example recording from human cortex during a seizure (see *Methods: Human Subject Data*). Visual inspection of the LFP data (Figure 11A) reveals the emergence of large amplitude voltage fluctuations during the approximately 80 s seizure. To analyze the CFC in these data, we separate this signal into 20 s segments with 10 s overlap, and analyze each segment using the proposed model framework. While little evidence of CFC appears before the seizure (Figure 11B), during the seizure we find significant **R**_PAC_ and **R**_AAC_ values. We show an example 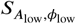 surface, and visualizations of this surface at small and large *A*_low_ values, in Figure 12. Repeating this analysis with the modulation index (Figure 11C), we find qualitatively similar changes in the PAC over the duration of the recording. However, we note that differences do occur. For example, at the segment indicated by the asterisk in Figure 11B, we find large **R**_AAC_ and an increase in **R**_PAC_ relative to the prior 20 s time segment, while increases in PAC and AAC remain undetected by **MI**.

**Figure 11.**
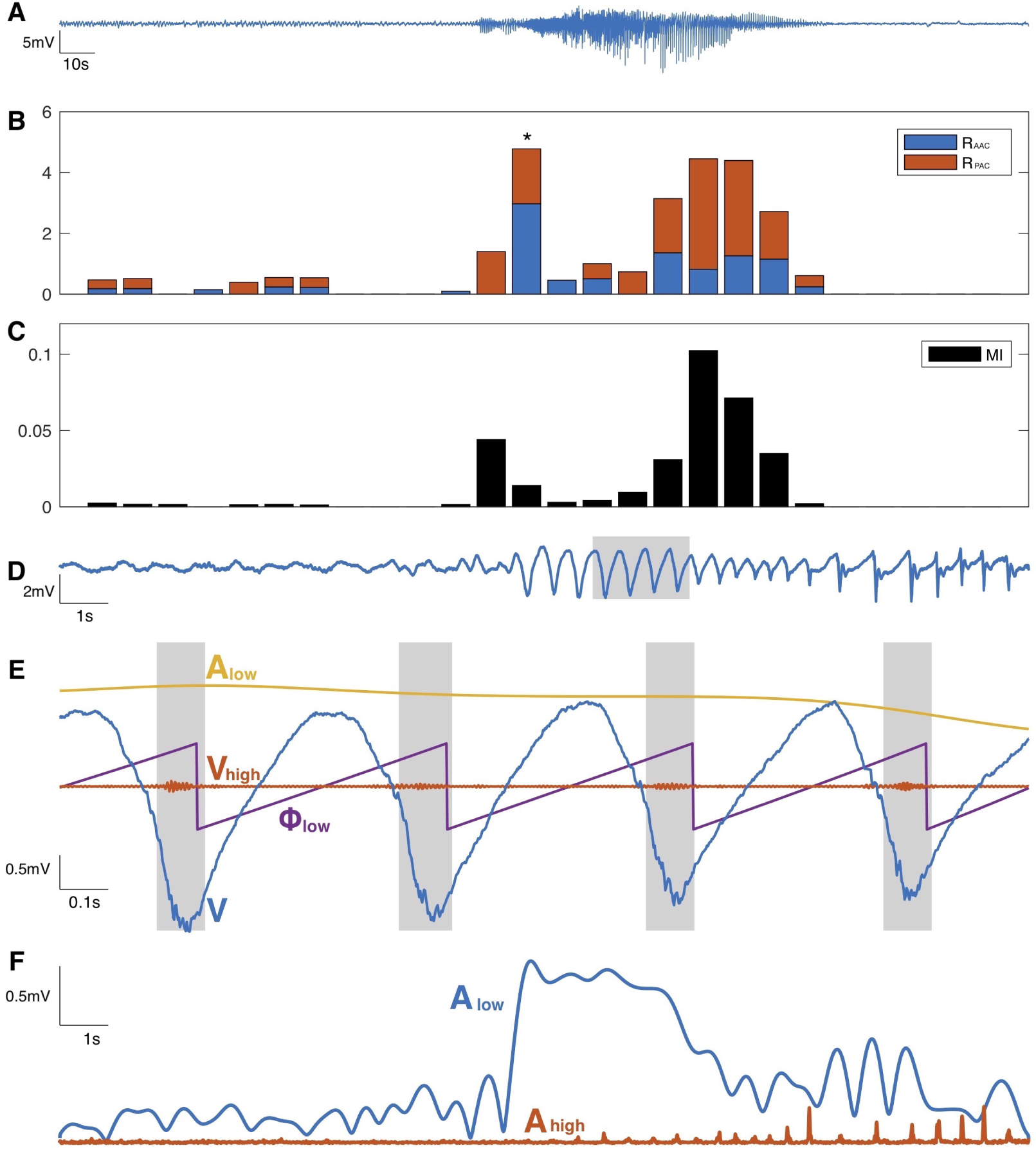
The CFC model detects cross-frequency coupling in an *in vivo* human recording. **(A)** Voltage recording from one MEA electrode over the course of seizure. **(B,C) R**_PAC_ and **R**_AAC_ values (B), and **MI** values (C), for 20 s segments of the trace. **(D)** Voltage trace corresponding to time segment (*) from (B). The gray-shaded time interval is analyzed in (E). **(E)** A subinterval of the voltage data (blue), along with *V*_high_ (red), *A*_low_ (yellow), and *ϕ*_low_ (purple). Grey shaded areas indicate increases in *V*_high_ when *ϕ*_low_ = 0. **(F)** *A*_low_ (blue) and *A*_high_ (red) for the 20 s segment in (D). *A*_high_ increases with *A*_low_ over time.

**Figure 12.**
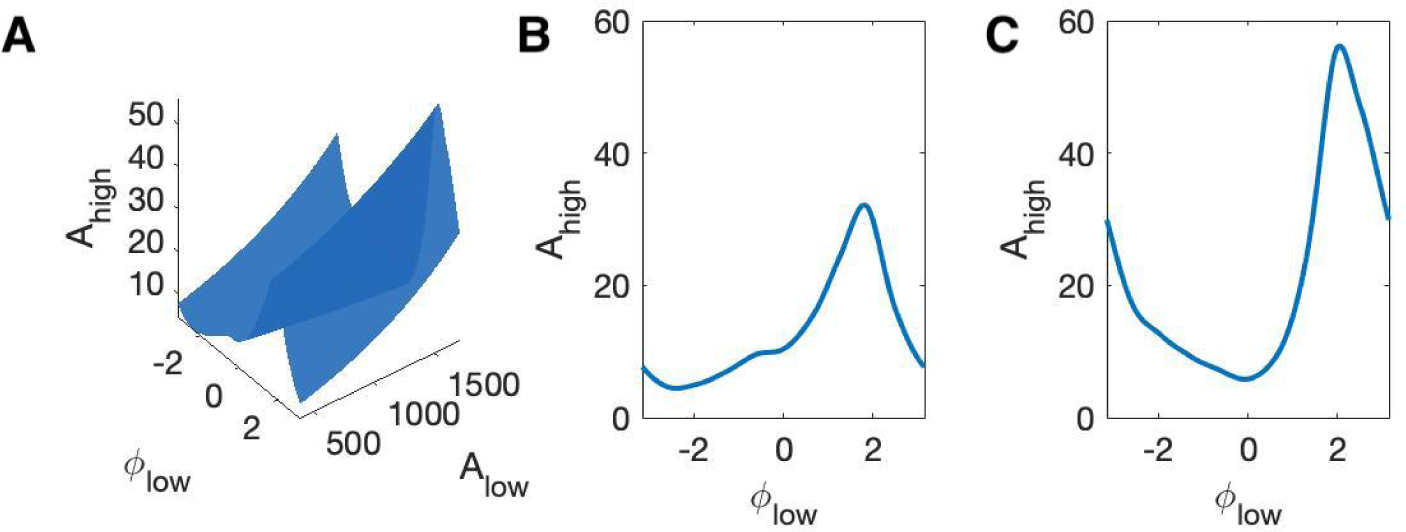
The 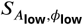 surface shows how PAC changes with the low frequency amplitude and phase during an interval of human seizure. **(A)** The full model surface (blue) in the (*ϕ*_low_, *A*_low_, *A*_high_) space, and components of that surface when **(B)** *A*_low_ is small, and **(C)** *A*_low_ is large.

To further investigate these results, we select a 20 s segment with significant and large **R**_PAC_ and **R**_AAC_ values to examine (Figure 11D). Visual inspection reveals the occurrence of large amplitude, low frequency oscillations and small amplitude, high frequency oscillations. To examine the detected coupling in even more detail, we isolate a 2 s segment (Figure 11E), and display the signal *V*, the high frequency signal *V*_high_, the low frequency phase *ϕ*_low_, and the low frequency amplitude *A*_low_. We observe that when *ϕ*_low_ is near *π* (gray bars in Figure 11E), *A*_high_ increases, consistent with the presence of PAC and a significant value of **R**_PAC_. Examining the low frequency amplitude *A*_low_ and high frequency amplitude *A*_high_ over the same 20 s segment (Figure 11F), we find that *A*_low_ and *A*_high_ increase together, consistent the presence of AAC and a significant value of **R**_AAC_.

### Application to *in vivo* rodent data

As a second example to illustrate the performance of the new method, we consider LFP recordings from from the infralimbic cortex (IL) and basolateral amygdala (BLA) of an outbred Long-Evans rat before and after the delivery of an experimental electrical stimulation intervention described in [5]. Eight microwires in each region, referenced as bipolar pairs, sampled the LFP at 30 kHz, and electrical stimulation was delivered to change inter-regional coupling (see [5] for a detailed description of the experiment). Here we examine how cross-frequency coupling between low frequency (5-8 Hz) IL signals and high frequency (70-110 Hz) BLA signals changes from the pre-stimulation to the post-stimulation condition. To do so, we filter the data *V* into low and high frequency signals (see *Methods*), and compute the **MI, R**_PAC_ and **R**_AAC_ between each possible BLA-IL pairing, sixteen in total.

We find three separate BLA-IL pairings where **R**_PAC_ reports no significant PAC pre- or post-stimulation, but **MI** reports significant coupling post-stimulation. Investigating further, we note that in all three cases, the amplitude of the low frequency IL signal increases from pre- to post-stimulation, and **R**_AAC_, the measure of amplitude-amplitude coupling, increases from pre- to post-stimulation. These observations are consistent with the simulations in *Results: The proposed method is less affected by fluctuations in low-frequency amplitude and AAC*, in which we showed that increases in the low frequency amplitude and AAC produced increases in **MI**, although the PAC remained fixed. We therefore propose that, consistent with these simulation results, the increase in **MI** observed in these data may result from changes in the low frequency amplitude and AAC, not in PAC.

### Using the flexibility of GLMs to improve detection of phase-amplitude coupling *in vivo*

One advantage of the proposed framework is its flexibility: covariates are easily added to the generalized linear model and tested for significance. For example, we could include covariates for trial, sex, and stimulus parameters and explore their effects on PAC, AAC, or both.

Here, we illustrate this flexibility through continued analysis of the rodent data. We select a single electrode recording from these data, and hypothesize that the condition, either pre-stimulation or post-stimulation, affects the coupling. To incorporate this new covariate into the framework, we consider the concatenated voltage recordings from the pre-stimulation condition *V*_pre_ and the post-stimulation condition *V*_post_:

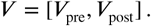

From *V*, we obtain the corresponding high frequency signal *V*_high_ and low frequency signal *V*_low_, and subsequently the high frequency amplitude *A*_high_, low frequency phase *ϕ*_low_, and low frequency amplitude *A*_low_. We use these data to generate two new models:

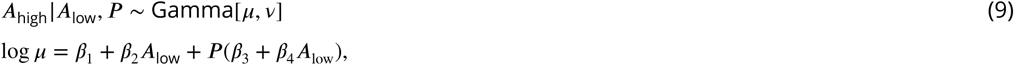

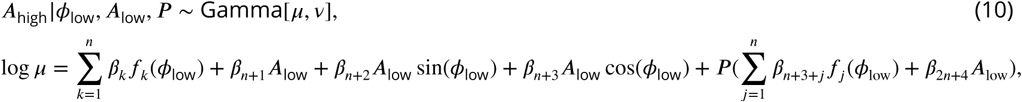

where *P* is an indicator function specifying whether the signal is in the *pre-stimulation* (*P* = 0) or *post-stimulation* (*P* = 1) condition. The effect of the indicator function is to add new terms to the *A*_low_ and *A*_low_, *ϕ*_low_ models to include the effect of stimulus condition on the high frequency amplitude. The model in Equation 9 now includes the effect of low frequency amplitude and condition on high frequency amplitude. The model in Equation 10 now includes the effect of low frequency amplitude, low frequency phase, and condition on high frequency amplitude.

To define **R**_PAC_, we construct a surface 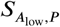 from the model in Equation 9 and a surface 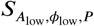 from the model in Equation 10 in the (*A*_low_, *ϕ*_low_, *A*_high_, *P*) space, assessing the models at the two values of *P*. We compute **R**_PAC_, which now measures PAC while accounting for changes in the low frequency amplitude and condition, as before:

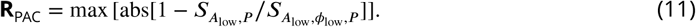

We find for the example rodent data an **R**_PAC_ value of 0.404 (*p* = 0.005). This provides strong evidence of PAC in the data, in at least one of the conditions.

This example illustrates the flexibility of the statistical modeling framework. Extending this framework is straightforward, and new extensions allow a common principled approach to test the impact of new predictors. Here we considered an indicator function that divides the data into two states (pre- and post-stimulation). More generally, an indicator function may specify a subset of conditions, during which a particular intervention or behavior occurred, and whether significant PAC occurs in any subset of conditions could then be tested. We note that the models are easily extended to account for multiple discrete predictors such as gender and participation in a drug trial, or for continuous predictors such as age and time since stimulus.

## Discussion

In this paper, we proposed a new method for measuring cross-frequency coupling that accounts for both phase-amplitude coupling and amplitude-amplitude coupling, along with a principled statistical modeling framework to assess the significance of this coupling. We have shown that this method effectively detects CFC, both as PAC and AAC, and is more sensitive to weak PAC obscured by or coupled to low-frequency amplitude fluctuations. Compared to an existing method, the modulation index [70], the newly proposed method more accurately detects scenarios in which PAC is coupled to the low-frequency amplitude. Finally, we applied this method to *in vivo* data to illustrate examples of PAC and AAC in real systems, and show how to extend the modeling framework to include a new covariate.

One of the most important features of the new method is an increased ability to detect weak PAC coupled to AAC. For example, when sparse PAC events occur only when the low frequency amplitude (*A*_low_) is large, the proposed method detects this coupling while other methods not accounting for *A*_low_ miss it. While PAC often occurs in neural data, and has been associated with numerous neurological functions [12, 31], the simultaneous occurrence of PAC and AAC is less well studied [53]. Here, we showed examples of simultaneous PAC and AAC recorded from human cortex during seizure, and we note that this phenomena has been simulated in other works [49].

While the exact mechanisms that support CFC are not well understood [31], the general mechanisms of low and high frequency rhythms have been proposed. Low frequency rhythms are associated with the aggregate activity of large neural populations and modulations of neuronal excitability [23, 76, 9], while high frequency rhythms provided a surrogate measure of neuronal spiking [58, 50, 27, 56, 83, 61, 60]. These two observations provide a physical interpretation for PAC: when a low frequency rhythm modulates the excitability of a neural population, we expect spiking to occur (i.e., an increase in *A*_high_) at a particular phase of the low frequency rhythm (*ϕ*_low_) when excitation is maximal. These notions also provide a physical interpretation for AAC: increases in *A*_low_ produce larger modulations in neural excitability, and therefore increased intervals of neuronal spiking (i.e., increases in *A*_high_). Alternatively, decreases in *A*_low_ reduce excitability and neuronal spiking (i.e., decreases in *A*_high_).

The function of concurrent PAC and AAC, both for healthy brain function and over the course of seizure as illustrated here, is not well understood. However, we note that over the course of seizure, particularly at termination, PAC and AAC are both present. As PAC occurs normally in healthy brain signals, for example during working memory, neuronal computation, communication, learning and emotion [71, 33, 12, 21, 38, 46, 37, 36, 66], these preliminary results may suggest a pathological aspect of strong AAC occurring concurrently with PAC.

Proposed functions of PAC include multi-item encoding, long-distance communication, and sensory parsing [31]. Each of these functions takes advantage of the low frequency phase, encoding different objects or pieces of information in distinct phase intervals of *ϕ*_low_. PAC can be interpreted as a type of focused attention; *A*_high_ modulation occurring only in a particular interval of *ϕ*_low_ organizes neural activity - and presumably information - into discrete packets of time. Similarly, a proposed function of AAC is to encode the number of represented items, or the amount of information encoded in the modulated signal [31]. A pathological increase in AAC may support the transmission of more information than is needed, overloading the communication of relevant information with irrelevant noise. The attention-based function of PAC, i.e. having reduced high frequency amplitude at phases not containing the targeted information, may be lost if the amplitude of the high frequency oscillation is increased across wide intervals of low frequency phase.

Like all measures of CFC, the proposed method possesses specific limitations. We discuss four limitations here. First, the choice of spline basis to represent the low frequency phase may be inaccurate, for example if the PAC changes rapidly with *ϕ*_low_. Second, the value of **R**_AAC_ depends on the range of *A*_low_ observed. This is due to the linear relationship between *A*_low_ and *A*_high_ in the *A*_low_ model, which causes the maximum distance between the surfaces 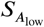 and 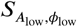 to occur at the largest or smallest value of *A*_low_. To mitigate the impact of extreme *A*_low_ values on **R**_AAC_, we evaluate the surfaces 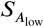 and 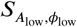 over the 5^th^ to 95^th^ quantiles of *A*_low_. We note that an alternative metric of AAC could instead evaluate the slope of the 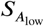 surface; to maintain consistency of the PAC and AAC measures, we chose not to implement this alternative measure here. Third, the frequency bands for *V*_high_ and *V*_low_ must be established before **R** values are calculated. Hence, if the wrong frequency bands are chosen, coupling may be missed. It is possible, though computationally expensive, to scan over all reasonable frequency bands for both *V*_high_ and *V*_low_, calculating **R** values for each frequency band pair. Fourth, we note that the proposed modeling framework assumes appropriate filtering of the data into high and low frequency bands. This filtering step is a fundamental component of CFC analysis, and incorrect filtering may produce spurious or misinterpreted results [3, 63, 41]. While the modeling framework proposed here does not directly account for artifacts introduced by filtering, additional predictors (e.g., detections of sharp changes in the unfiltered data) in the model may help mitigate these filtering effects.

The proposed method can easily be extended by inclusion of additional predictors in the GLM. Polynomial *A*_low_ predictors, rather than the current linear *A*_low_ predictors, may better capture the relationship between *A*_low_ and *A*_high_. One could also include different types of covariates, for example classes of drugs administered to a patient, or time since an administered stimulus during an experiment. To capture more complex relationships between the predictors (*A*_low_, *ϕ*_low_) and *A*_high_, the GLM could be replaced by a more general form of Generalized Additive Model (GAM). Choosing GAMs would remove the restriction that the conditional mean *A*_high_ must be linear in each of the model parameters (which would allow us to estimate knot locations directly from the data, for example), at the cost of greater computational time to estimate these parameters. The code developed to implement the method is flexible and modular, which facilitates modifications and extensions motivated by the particular data analysis scenario. This modular code, available at https://github.com/Eden-Kramer-Lab/GLM-CFC, also allows the user to change latent assumptions, such as choice of frequency bands and filtering method. The code is freely available for reuse and further development.

Rhythms, and particularly the interactions of different frequency rhythms, are an important component for a complete understanding of neural activity. While the mechanisms and functions of some rhythms are well understood, how and why rhythms interact remains uncertain. A first step in addressing these uncertainties is the application of appropriate data analysis tools. Here we provide a new tool to measure coupling between different brain rhythms: the method utilizes a statistical modeling framework that is flexible and captures subtle differences in cross-frequency coupling. We hope that this method will better enable practicing neuroscientists to measure and relate brain rhythms, and ultimately better understand brain function and interactions.

## Acknowledgements

This work was supported in part by the National Science Foundation Award #1451384, in part by R21 MH109722, and in part by the National Science Foundation (NSF) under a Graduate Research Fellowship.

